# An Advanced Mobile Laboratory to enable field-based microbial ecology and cell biology across scales

**DOI:** 10.64898/2026.02.23.707475

**Authors:** N Leisch, S Baars, T Beavis, P Bertucci, C Bhickta, M Bonadonna, CM Brannon, L Burgués-Palau, P Cherek, F Chevalier, J Decelle, M Demulder, G Dey, O Dudin, E Duke, BD Engel, E Flaum, S Flori, B Gallet, P Guichard, A Halavatyi, V Hamel, M Jacobovitz, C Juery, M Laporte, S Mattei, F Mikus, K Mocaer, K Moog, M Olivetta, M Pavie, R Pepperkok, A Perez-Boerema, L Planat, M Prakash, EW Pyle, CR Rhodes, I Romero-Brey, P Ronchi, H Rosa, A Rubio Ramos, C Saint-Donat, Y Schwab, H Shah, AM Steyer, A Svetlove, G Toullec, F Vincent, T Wiegand, W Wietrzynski, D Yee, S Zwahlen

**Affiliations:** Mobile Laboratories Services, European Molecular Biology Laboratory Heidelberg, Germany; Biozentrum, University of Basel. Switzerland; Developmental Biology, European Molecular Biology Laboratory, Heidelberg, Germany; Cell Biology and Biophysics, European Molecular Biology Laboratory Heidelberg, Germany; Faculty of Biology, Heidelberg University, Heidelberg, Germany; Department of Biology, Stanford University, USA; Cell and Plant Physiology Laboratory, CNRS, CEA, INRAE, IRIG, Université Grenoble Alpes, Grenoble, France; Department of Biochemistry, Faculty of Sciences, University of Geneva, Geneva, Switzerland; Biological X-ray Imaging, European Molecular Biology Laboratory Hamburg, Germany; Graduate Program in Biophysics, Stanford University, Stanford, USA; Department of Bioengineering, Stanford University, Stanford, USA; Université Grenoble Alpes, CEA, CNRS, IBS, Grenoble, France; Department of Molecular and Cellular Biology, Faculty of Sciences, University of Geneva, Switzerland; Advanced Light Microscopy Facility, European Molecular Biology Laboratory Heidelberg, Germany; Université Claude Bernard Lyon 1, MeLiS, CNRS UMR95284, INSERM U1314, Lyon, France; Molecular Systems Biology. European Molecular Biology Laboratory Heidelberg, Germany; EMBL Imaging Centre, European Molecular Biology Laboratory Heidelberg, Germany; Centre for Organismal Studies, Heidelberg University, Heidelberg Germany; Bioengineering, Stanford University, USA; Electron Microscopy Core Facility, European Molecular Biology Laboratory Heidelberg, Germany

## Abstract

Microbial biodiversity is central to ecosystem function, yet mechanistic insights into the cell biology of environmental organisms remain limited. The underlying challenges are twofold: most microbes remain uncultivable, and a persistent gap exists between field sampling and laboratory analyses. Here, we introduce the Advanced Mobile Laboratory (AML), a field-deployable platform that integrates confocal microscopy, image-enabled cell sorting, and cryo-preparation for expansion and electron microscopy. This setup enables immediate, standardized processing and analysis of environmental communities directly at the sampling site. We demonstrate its capability using marine eukaryotic plankton, showing how the AML enables multiscale investigations, from live imaging of natural communities to enabling ultrastructural and single-cell omics analyses, while minimizing sample degradation and enabling on-site experimentation. By bringing high-end sample preparation and analytical capacity into the field, the AML enables studying life in its natural context to mechanistically understand life’s diversity in the environment.

## Introduction

The current era represents a golden age of biodiversity discovery, particularly for microbial life ranging from viruses to protists ^1–5^. However, this biodiversity is globally threatened ^6–8^, and consequently, there is an urgent need to advance a mechanistic understanding of cellular life in its environmental context, to expand efforts in studying biodiversity, and to understand the adaptive capabilities of species.

Field-based research has driven landmark advances in biology, from Darwin’s voyage on the HMS *Beagle*, which led to his formulation of the theory of evolution, to Chatton’s eukaryote-prokaryote distinction and the foundational contribution of the Challenger expedition to oceanography ^9–11^. While science has shifted towards large centralized labs over the last century, field work is central to understanding earth’s changing biodiversity, biogeochemical cycles, and the interconnectedness of earth’s habitats ^12–14,reviewed in 15^. In parallel, technological advances have transformed our understanding of these biological processes and how we study microbial communities. Genetic sequencing has revealed the taxonomic and functional diversity in both environmental and host-associated microbial communities (e.g. ^16,17^), while diverse microscopy approaches capture aspects of microbial structure and organization ^18–23,reviewed in 24^. Yet each approach has limitations, high-throughput sequencing lacks spatial resolution and microscopy methods involve trade-offs between resolution, throughput, and sample preparation.

Despite these major advances in sequencing and imaging, core challenges persist in microbial research. The vast majority of microbial life remains uncultivable using standard laboratory techniques, which skews our understanding toward a few model taxa and limits our ability to recreate and study complex ecologies in the lab ^25^. Moreover, while we can rapidly profile the taxonomic and functional diversity of microbial communities across ecosystems (e.g. ^1,5,26^), we still lack the cellular and physiological context needed to connect the genotype to the phenotype of these organisms and to understand their interactions with and within their native habitat. Visually capturing biology that spans communities, cellular interactions, subcellular structures and protein localization, requires crossing scales from light and fluorescence microscopy to electron and x-ray imaging ^27–29^.

Bridging these gaps requires integrating high-resolution imaging, cultivation, and molecular approaches directly at the point of sampling in the environment. Field-deployed technologies have repeatedly revealed new biological insights (e.g.,^30^), demonstrating the power of on-site analysis. Here, as a proof of concept, we built the Advanced Mobile Lab (AML), outfitted with confocal microscopy, cell sorting, and cryo-preparation for electron microscopy, to enable the rapid transition from environmental sampling to imaging and analysis. By minimizing the delay between collection and processing, the AML mitigates some of the logistical and methodological constraints that are intrinsic to fieldwork. In addition, the AML supports controlled experiments and data acquisition on site, which typically are restricted to cultured systems, thereby paving the way for a stronger link between field- and lab-based research approaches. We demonstrate the opportunities the AML enables for marine plankton biodiversity research, which form the base of ocean food webs and regulate climate via carbon cycling ^reviewed in 15,26,reviewed in 31^, with a focus on micro-eukaryotes, across scales from proteins to complex communities.

## Results and discussion

### The AML - a modern molecular lab on wheels

The key considerations behind the inception of the AML were threefold. First, bringing the lab into the field enables direct experimentation with and analysis of live organisms, and provides controlled conditions for preserving samples directly in the field, maximizing sample quality. Second, deploying advanced analysis technologies directly on site enables complex workflows and iterative, adaptive sampling schemes. Finally, deploying the same set of instruments, protocols and trained application scientists to each sampling site provides an unprecedented degree of standardization and reproducibility.

The AML consists of a customised semi-trailer that houses a modern molecular laboratory (**Figure 1 A, Supplementary Note**). Along the length of the AML, two slideouts can be hydraulically extended to increase the interior work space to ~40m^2^ (**Figure 1 B,C**), accommodating 8-12 researchers simultaneously.

**Figure 1.**
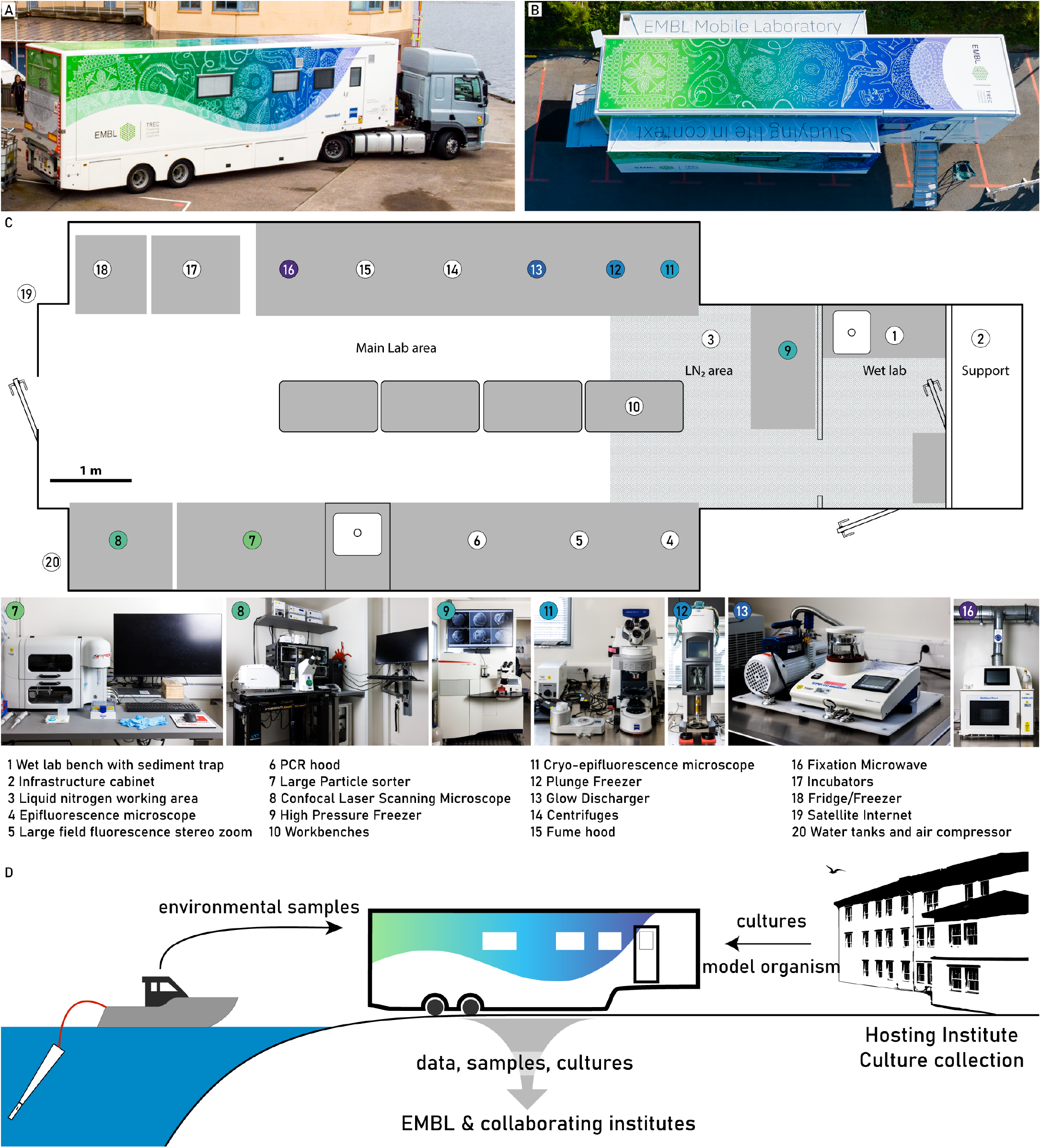
Overview of the Advanced Mobile Lab, its technologies and the sample and data workflow. The AML in its travel configuration (**A**) and fully deployed, with the slide-outs extended (**B**). (**C**) shows a schematic overview of the lab, indicating the key areas and instruments (bottom row, and highlighted in color) and the overall instrument distribution. (**D**) Samples are retrieved from the environment close to the lab or alternatively from the host institute or a culture collection nearby. Data and samples generated in the AML are distributed to EMBL and the collaborating institutes before departure.

The lab is staffed by four experts that operate the key technologies: 1) a large particle sorter with imaging capabilities that enables rapid isolation of target organisms from diverse environmental samples, over a wide size range. 2) A broad range of light microscopy equipment including simple binoculars, a light and fluorescence microscope, a large-field fluorescence binocular and a confocal laser scanning microscope. 3) A microwave-assisted tissue processor, a high-pressure freezer and a plunge freezer for electron microscopy sample preparation. Additionally, a fluorescence microscope with a cryo-stage is available for on-site screening and quality control of the vitrified specimens.

Environmental samples are brought directly into the wet lab area for further processing (e.g. filtration of seawater, sediment extraction, etc), and to keep corrosive elements away from the sensitive instruments (**Figure 1 C,D**). The main laboratory has a dedicated area for work with liquid nitrogen, which is equipped with a direct link to an external LN_2_ dewar. Furthermore, the AML is outfitted with all the necessary infrastructure for a modern molecular biology lab, which includes a fume hood, a PCR preparation hood, a dangerous goods cabinet, centrifuges, a fridge and a freezer. Multiple incubators are available to initially store samples at adequate light and temperature conditions, as well as enable incubations, cultivation or experimentation on site (**Figure 1 C**).

We designed transport safeguards for all the equipment to minimize vibrations and prevent damage during the transport, allowing most instruments to remain installed during transport with minimal disassembly. On-site setup involves disconnecting the trailer from the truck, stabilizing and levelling the AML with hydraulic supports, supplying power to the AML, and extending the slide-outs. This process can be done in as little as 20 minutes. Preparing most machines for operation takes an additional 4 to 6 hours, with some requiring more time (e.g., overnight stabilisation for the lasers).

The AML is designed to be as modular as possible to easily accommodate changes or replacement of the instruments as well as project specific requirements. Furthermore, we used the AML to field-test new technologies and prototypes throughout the deployment. For such technology trials, the AML provides the necessary support of a sophisticated laboratory in close proximity to the biology.

### The AML in the context of the Traversing European Coastlines expedition

The AML was initially built to support the Traversing European Coastlines expedition (TREC) expedition led by EMBL, in collaboration with EMBRC and the Tara Oceans Foundation, which took place from 2023-2024. Over 18 months, researchers collected comprehensive samples across 21 countries to investigate critical ecological issues, including biodiversity loss, pollution, antibiotic resistance, and the anthropogenic impact on habitats (https://www.embl.org/about/info/trec/). Altogether, the AML supported over 70 researchers across eight locations. The AML enabled us to reproducibly apply advanced assays in standardised and systematic ways across different ecosystems (**Figure 1 D**).

### From screening to 3D - Confocal analysis of live plankton communities

Confocal microscopy has been routinely used in the marine sciences, enabling researchers to image subcellular structures in plankton in 2D and 3D, relying on a combination of dyes and naturally fluorescent pigments to identify cells ^18,32^. We systematically screened all freshly collected plankton samples, which informed downstream experiments conducted on that day. Additionally, these images serve as a snapshot of the biodiversity and abundance in these samples and are publicly available in the framework of the BIOcean5D EU project (**Figure 2 A**) ^33,34^. Furthermore, the confocal microscope enables recording 3D data of selected specimens to better understand taxonomy, cell states and interactions (**Figure 2 B-F**). To enable this in a high-throughput manner, we applied feedback microscopy ^35,36^ on living organisms of interest. As an example, we targeted organisms containing the light-harvesting pigment phycoerythrin, but this workflow can be extended to other features (**Supplementary Figure 1**). Additionally, imaging live samples avoids fixation bias, enables behavioral observation (**Supplementary Movie 1**), and can be combined with dyes to probe cellular physiology and ultimately represent planktonic communities closer to their native physiological state.

**Figure 2.**
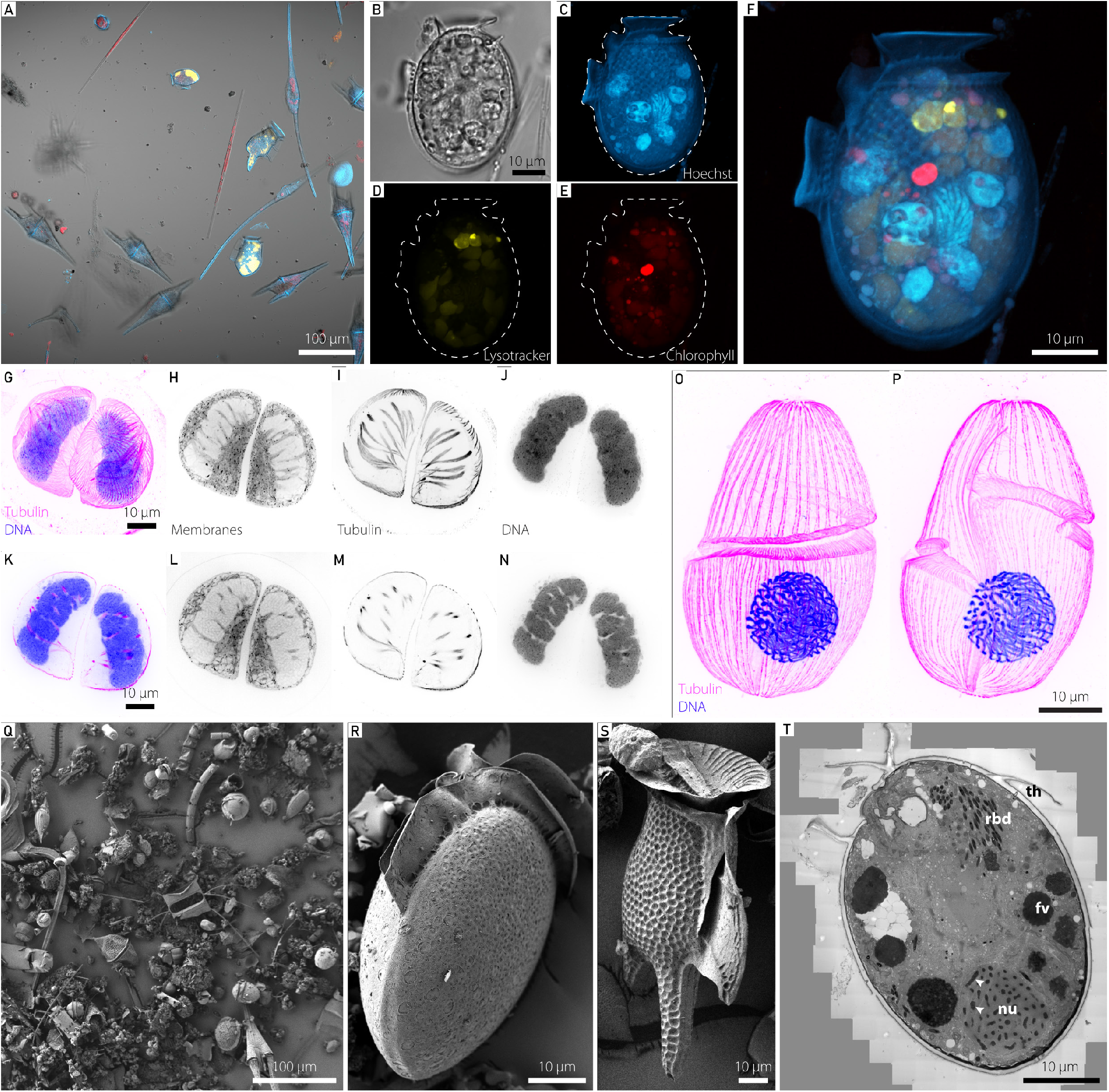
From diverse live plankton communities to subcellular morphology. **Top row (A)** Overview image of a typical environmental plankton sample imaged by CLSM (red: chlorophyll autofluorescence, yellow: phycoerythrin autofluorescence, blue: Hoechst staining, grey: transmitted light). **(B-F)** Maximum intensity projection of a single Dinophysis sp. cell imaged as a high-resolution z-stack **(B)** transmitted light, **(C)** Hoechst staining in blue, **(D)** Lysotracker-green staining in yellow, (E) chlorophyll autofluorescence in red, **(F)** overlay of C-E. **Second row (G-P)** Ultrastructure Expansion Microscopy of a plunge-frozen dinoflagellate. Staining for tubulin (magenta) and DNA (blue) reveals their cortical microtubule network and highly condensed chromosomes, and membrane staining provides cellular context. **(G)** shows the maximum intensity projection of tubulin and DNA stain. **(H-J)** show the medial maximum intensity projection of the membrane stain, tubulin stain, and DNA stain, respectively. **(K-N)** show a single confocal slice each in the same order as **(G-J). (O**,**P)** show a maximum intensity projection through the dorsal **(O)** and ventral **(P)** half of another dinoflagellate, possibly *Gyrodinium* sp., cell. Scale bars were adjusted for the expansion factor. **Third row (Q-S)** Scanning electron microscopic analysis, highlighting the diverse community in **(Q)**, individual *Dinophysis* cells in **(R**,**S)** and transmission electron microscopy analysis providing subcellular analysis of a single *Dinophysis* sp. cell in **(T)** (high-resolution TEM image shown in Supplementary Figure 2). Th-theca, fv-food vacuole, rbd-rhabdosomes, nu-nucleus, white arrowheads-chromosomes.

### Democratizing expansion microscopy for environmental cell biology

While confocal approaches are becoming increasingly standardized and automated for high-throughput imaging, especially of single celled organisms, it can quickly be limited by the resolution and the lack of epitope accessibility for dyes and antibodies. To address these limitations, expansion microscopy (ExM) is a recently developed super-resolution method that enables high-resolution fluorescence imaging of fixed cells ^37,38^. The core principle of ExM involves embedding fixed samples into swellable hydrogels, followed by the disruption of protein links, either through digestion steps or denaturation, and finally the physical expansion of the sample.

In particular, the ultrastructure expansion microscopy (U-ExM) protocol is compatible with a broad range of protists, mammalian cells, and yeast species ^39–46^. This significantly minimizes the need to adjust protocols for dyes, stains, and antibodies across different organisms and enables us to, as in one recent example, study cytoskeletal diversity throughout the tree of life ^47^. Thanks to the AML-enabled deployment of instruments for the vitrification of specimens on site, we applied cryo-ExM ^43^ to concentrated fresh plankton samples. They were cryo-preserved either with a high-pressure freezer ^48^ or a customized plunge freezer designed for microscopy coverslips ^43^. Vitrified samples can often exhibit superior epitope and ultrastructural preservation, particularly for cytoskeletal proteins and membranes as shown here for a dinoflagellate cell from the environmental sample (**Figure 2 G-P**) and ^49^ for diatoms. The broad applicability, resolution gain and high-throughput makes expansion microscopy an ideal approach to complement light and electron microscopy and bridge the resolution gap between these two.

### Investigating environmental plankton ultrastructure using 2D electron microscopy

Electron microscopy has been essential for plankton research, it provides subcellular information, aids taxonomic efforts ^50,51^, and enables the detection and morphometric quantification of novel structures and organelles ^52,53^.

We applied both microwave assisted fixation for Scanning Electron Microscopy (SEM) analyses and cryo-fixation-based methods for the Transmission Electron Microscopy (TEM) analyses of plankton communities (**Figure 2 Q-T**). This approach allowed us to analyse the community composition (**Figure 2 Q-S**) and the subcellular ultrastructure, here of the dinoflagellate *Dinophysis* sp. (**Figure 2 T**). Especially for subcellular ultrastructure, cryo-fixation is crucial and enables the visualisation of cellular ultrastructures in their near-native state ^54,55^.

### Towards environmental structural biology

One of the most challenging frontiers when working with environmental microbial communities is leveraging cryo-electron tomography (cryo-ET) to access the diversity and function of macromolecules and proteins within the cellular context ^56,57^. For example, cryo-ET has been key to understanding pyrenoids, specialized CO_2_-concentrating compartments that enable algae to perform efficient carbon fixation ^58^. When applied to diatoms, which fix about 20% of global CO_2_, cryo-ET revealed that their pyrenoids are surrounded by a protein sheath called the PyShell, which is crucial for efficient carbon fixation as well as structural integrity of the pyrenoid ^59,60^. With rapid ongoing developments in both instrumentation and data processing, cryo-ET is spearheading the new field of visual proteomics, which promises to reveal molecular inventories of the native cellular environment ^27^.

Here, we used plunge freezing, which can vitrify cells smaller than 10 µm, to collect diverse planktonic species on grids (**Figure 3 A-K**). We used two workflows to target either dinoflagellate (**Figure 3 A-D**) or diatom cells (**Figure 3 E-K**). Using cryo-SEM we identified abundant ovoid-shaped cells of similar size (**Figure 3 A**) to target for Focused Ion Beam (FIB) milling (**Figure 3 B**). We acquired, reconstructed, and segmented tomograms from the milled lamellae, confirming that these cells were dinoflagellates, characterized by their distinctive condensed chromosomes 61,62 and crystalline structures surrounding the chloroplast (Jantschke et al., 2019) (**Figure 3 C and D**). In parallel, we used cryo-light microscopy to survey the cell population and identify a Pseudo-nitzschia sp. cell on the frozen grid (**Figure 3 E and F**). After milling cryo-lamellae (**Figure 3 G**), acquiring tomograms, and segmenting the organelles (**Figure 3 H and I**), we performed subtomogram averaging to further characterize the structures of cytosolic ribosomes (**Figure 3 J-K**). Although the attained resolution was moderate due to limited particle number in this environmental sample, clear structural features were resolved, as well as indications of polyribosome organization. Expanding cryo-ET methodology to specifically target members of environmental communities offers exciting opportunities to study the native molecular architecture of these organisms in the context of external factors such as seasonal changes, light and nutrient availability, or in the case of plankton, blooming events. So far, cryo-ET has mostly focused on model species, but applying this approach to environmental samples will allow testing of evolutionary and functional hypotheses developed in controlled settings. This will make it possible to link organelle and protein-level adaptations to species diversity, bridging the gap between laboratory models and the complexity of natural ecosystems.

**Figure 3.**
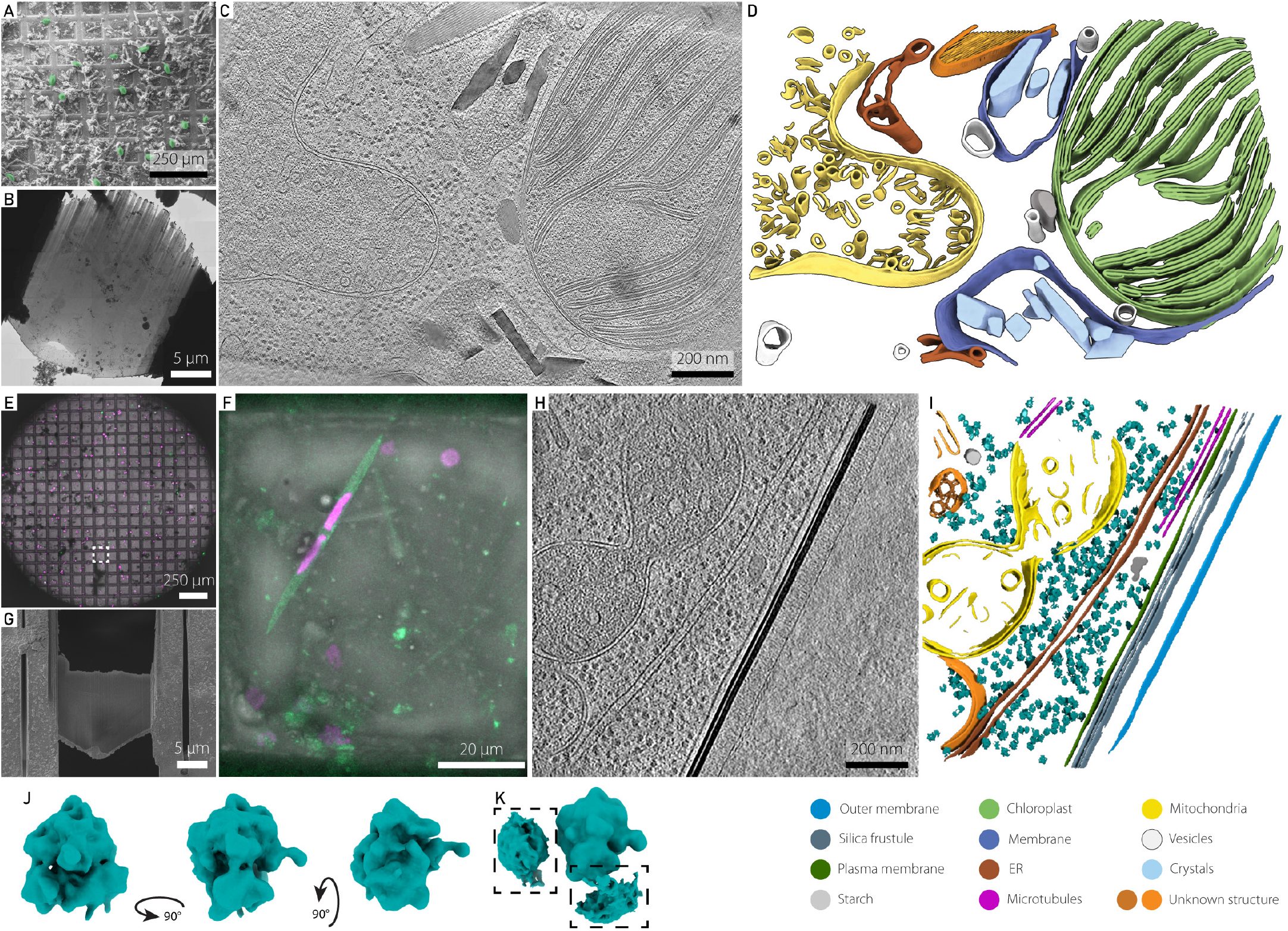
Cryo-electron tomography workflows for environmental samples. **Top row (A-D)** Cryo-electron tomography workflow from left to right showing **(A)** overview imaging in cryo-SEM to identify targets within a plunge frozen environmental sample (highlighted in green pseudocolor), **(B)** cryo-FIB lamella preparation of a target cell, followed by cryo-ET data acquisition, **(C)** tomogram reconstruction, and **(D)** 3D segmentation of corresponding volume. **Bottom row (E-K)** Correlative cryo-light and -electron tomography workflow from left to right showing **(E)** the overview of the frozen grid in the cryo-confocal, with transmitted and reflected light as well as autofluorescence in 488 nm (green) and 567 nm (magenta). **(F)** shows a maximum intensity projection of a single grid square, corresponding to the dashed line in **E. (G)** Scanning electron microscopy image of milled lamella in the region highlighted by dashed line in **E. (H)** A slice through the reconstructed tomogram and the corresponding 3D segmentation. **(J)** Isosurface representation of the ribosome subtomogram average (29 Å resolution), shown from three viewing angles. **(K)** Subtomogram average from **J** after denoising by M software, shown from two viewing angles at a lower isosurface threshold. Highlighted in boxes are densities corresponding to neighbouring ribosomes from particles in a polysome arrangement.

### I choose you - Capturing the subcellular phenotype of a target species from the environment

The above approaches collect and preserve the whole microbial comm unity from the environment. Therefore, the identification, isolation and analysis on a particular organism of interest across samples is very challenging and time-consuming, and can be achieved only *a posteriori* after sample preservation. Importantly, there is a substantial risk to miss the organism in the complex community. We therefore developed a specific workflow to adopt a reverse approach: starting with a freshly collected environmental sample, live and unlabelled cells of the same organism are rapidly isolated with a cell sorter and directly deposited in a custom sample holder to enable a swift transfer to the high pressure freezer and subsequent downstream analyses (**Figure 4**). This avoids manual sorting for individual cells, which can be arduous or even impossible for small cells, it reduces the crucial time between organism identification and preservation, and most importantly it improves the reproducibility and statistical power needed to capture the potential heterogeneity of the population in the environment. As a proof of concept, from an environmental planktonic community ranging from 20 to 200 µm in size, we sorted hundreds of cells which showed an autofluorescence signal for both chlorophyll and phycoerythrin and were morphologically similar to the dinoflagellate *Dinophysis* sp. (~62.8 µm in length). Within ~5 minutes of sorting, around 300 cells of *Dinophysis sp*. were isolated, with these cells representing ~1% of the environmental community (**Figure 4 A and B**). The cells were directly deposited into a custom holder for the high-pressure freezing carrier (**Figure 4 C**), which facilitated immediate cryofixation with a high-pressure freezer (**Figure 4 D**). Following cryo-fixation, the sorted cells were quality-controlled using a cryo-fluorescence microscope, with the chlorophyll autofluorescence shown in (**Figure 4 E**) indicating both high purity of the sorting and intact cells after the vitrification. After return to the respective research institutions, sorted and vitrified cells were cryo-substituted and embedded in a resin for subcellular investigation using volume electron microscopy (vEM) (**Figure 4 F**), specifically Focused Ion Beam Scanning Electron Microscopy (FIB-SEM). FIB-SEM imaging records isotropic datasets at nanometric resolution, and associated morphometric analyses bring new insights on single cell ultrastructure (Mocaer et al) or complex cell-cell interactions in the plankton ^46,54,55^, especially to disentangle the spatial organisation of the host and symbiont organelles and link with their functional roles. To target the vEM acquisition precisely on a cell, high throughput X-ray tomography (HiTT) ^63^ was performed on a resin block containing the sorted cells. HiTT provides an isotropic 3D volume of the resin block, from which we identified and segmented 14 cells for morphometric analysis and recorded the position of these cells in relation to topological landmarks of the resin block (**Figure 4 G, Supplementary Table 1**). These coordinates guide the downstream FIB-SEM imaging ^64^. Here, we acquired FIB-SEM data (a stack of 2270 images with isotropic 20 nm voxel size) and segmented multiple organelles from a single *Dinophysis sp*. cell (**Figure 4 H-J, Supplementary Video 2**). *Dinophysis* is an example of “third-hand kleptoplastidy”, stealing chloroplasts from its ciliate prey *Myrionect*a, which itself previously acquired its chloroplasts from cryptophyte microalgae ^65–67^. Based on the HiTT-derived morphometrics and the FIB-SEM data, we morphologically identified the cells as either *D. norvegica* or belonging to the *D. acuminata* species complex. Both are abundant in the Baltic Sea and have overlapping morphological features ^68^. Here, our 3D reconstruction of a single cell (**Figure 4 H-J**) revealed the structural organisation of chloroplasts. The 15 segmented chloroplasts ranged from 12.32 µm^3^ to 259.02 µm^3^ in volume (mean 73.35 ± 81 µm^3^), and contained multiple pyrenoids (**Figure 4 I and J, Supplementary Table 1**). Surrounding the chloroplasts, several starch grains were visible, reflecting storage of sugars derived from photosynthesis (**Figure 4 J**). The dinoflagellate nucleus had a volume of 2,452 µm^3^ and our 3D reconstruction revealed at least 267 chromosomes (mean 1.55 ± 0.99 µm^3^ each), occupying 17.07% of the nuclear volume. The number of chromosomes is relatively high and voluminous compared to other dinoflagellate species: ~113–119 in *Scrippsiella* ^69^, from 64 to 179 in *Brandtodinium* ^70^, 105 chromosomes in *Ensiculifera* ^55^, except for *Oxyrrhis marina* with ca. 400 chromosomes ^71^. The dataset also allowed for reconstruction of the peduncle (**Figure 4 I**), an organelle involved in the formation of an extensible tubular structure which enables nutrient acquisition without full engulfment of the prey cell ^72,73^ as well as structures like the enigmatic rod-shaped rhabdosomes, whose function might be related to prey capture or defense but is still unclear ^52^.

**Figure 4.**
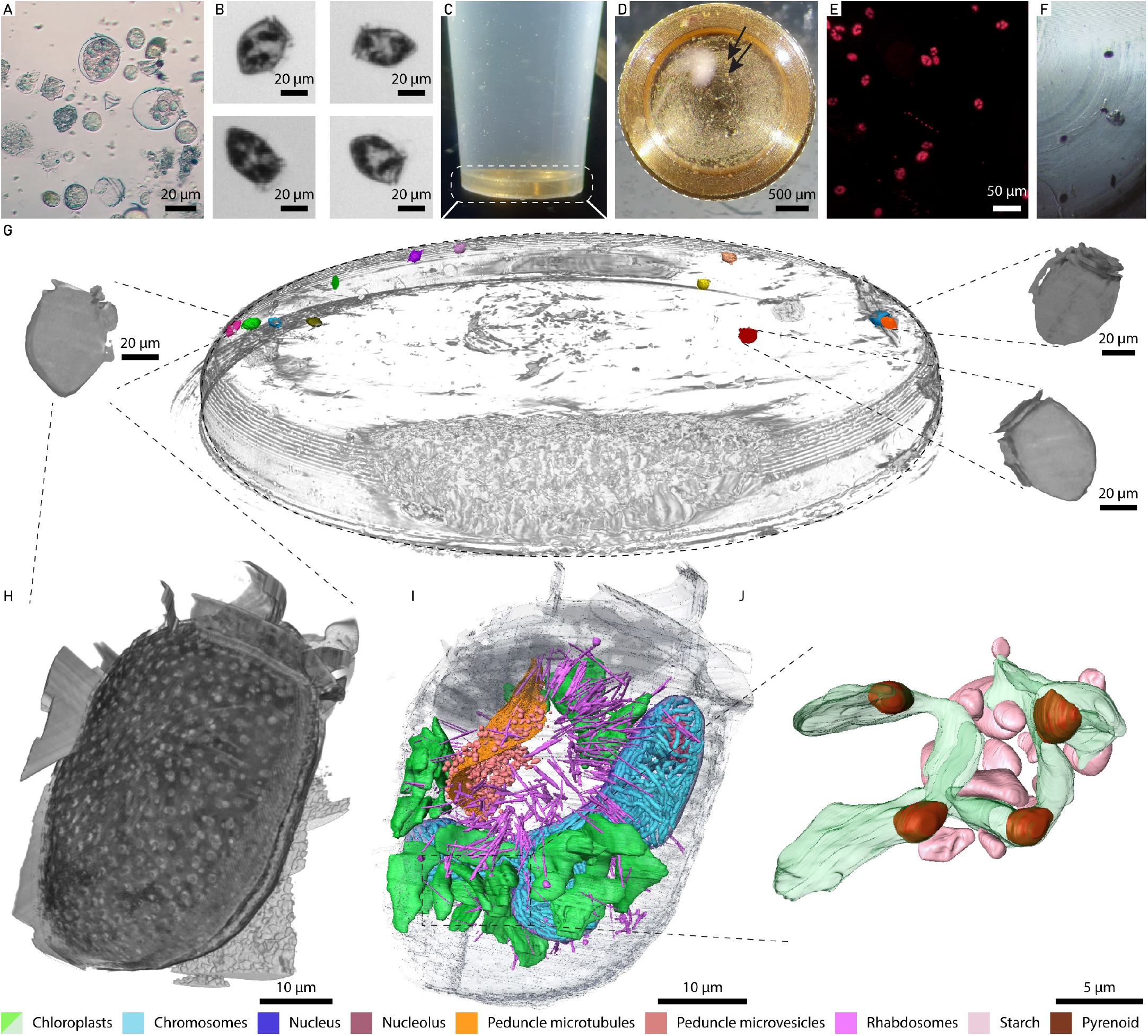
Targeted workflow to isolate a plankton species from the community and reveal its external and subcellular morphology. **(A)** The environmental plankton sample from which **(B)** the dinoflagellate *Dinophysis* sp. was sorted and transferred into a custom-made HPF carrier holder **(C). (D)** shows the quality control before freezing (arrows indicate single cells), **(E)** shows the chlorophyll fluorescence-based detection in the frozen carrier and **(F)** the individual cells embedded in the resin block after freeze substitution. **(G)** 3D-rendering of the HiTT-imaged whole block. All cells were identified within the block and segmented. Three segmented cells are shown in a close up isolated from the whole block. **(H-J)** A single *Dinophysis* sp. cell was imaged by FIB-SEM. A 3D rendering of the cell boundaries allowed assessment of the external cellulosic theca (grey), providing morphological cues for species identification **(H)**, and subsequent segmentation of the vEM data reveals subcellular features **(I and J). (J)** shows a zoom in on a single chloroplast volume, indicating the pyrenoids and the starch granules.

In summary, we have developed an original workflow to - without any pre-labeling steps - rapidly isolate a live organism of interest which occurs at a low abundance, from a freshly collected, complex environmental community, cryo-preserve, and analyze with a range of imaging modalities to gain nanoscale phenotypic information in *natura*. This workflow allows one to reveal not only the complex cellular architecture of the isolated organisms in a given environmental condition and location, but also to disentangle cryptic symbiotic associations with other microbial partners or stolen organelles.

### Bridging lab and field research

A key step towards understanding biodiversity is the organism-centric integration of molecular and cellular data types as well as conducting process-oriented studies such as time series and incubation experiments. These approaches are often restricted to model organisms that can be grown at high concentration and high purity in the lab’s controlled conditions. Here, the AML offers the ability to integrate a vast array of sensitive methods on non-model systems, and contributes to building the necessary bridge between lab- and field-based research approaches.

We identified *Sundstroemia sp*. as the target organism of our integrative approach, in one of our environmental samples. This diatom is a silicified phytoplankton, previously classified as *Rhizosolenia sp*., and belongs to the family of bloom-forming diatoms. Following image-enabled cell sorting to enrich and purify the target species (**Figure 5 A-C**), we performed three categories of assays in parallel: (i) process-oriented assays that require live imaging, (ii) time-sensitive assays needing immediate preservation to capture highly dynamic properties, and (iii) biodiversity assays demanding high sensitivity and minimal contamination.

**Figure 5.**
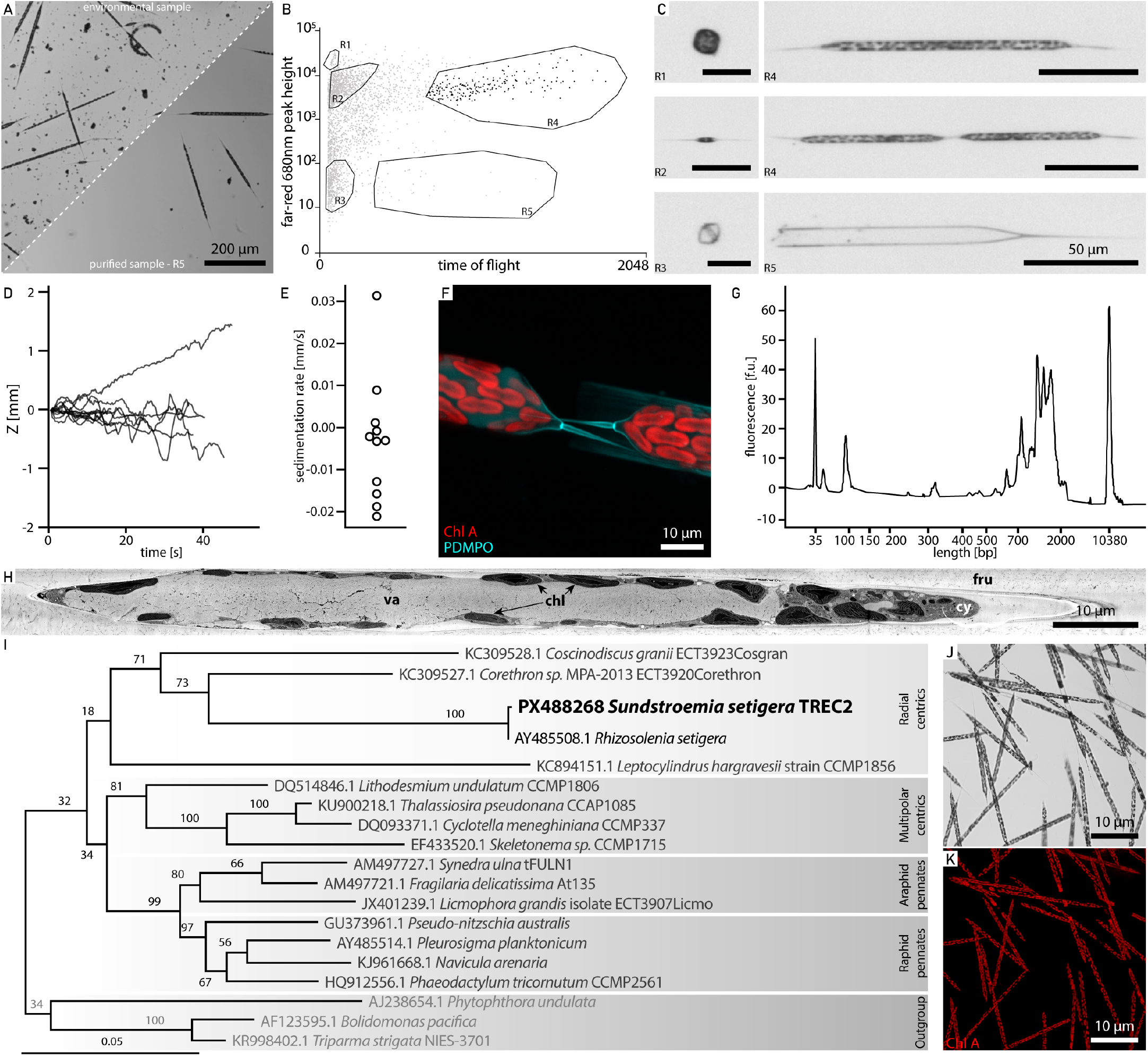
Multi-modal analysis of environmental marine microbes. **(A)** Environmental sample containing the key player *Sundstroemia setigera* before (top half) and after (bottom half) cell sorting. **(B)** Gating strategy to purify *S. setigera* and **(C)** corresponding images, where the labels R1-R5 in **(B)** correspond to the images in **(C)**. **(D)** Sinking distance of *S. setigera*, through time and **(E)** computed sedimentation rate. **(F)** Confocal image of incorporated silica after 72 hours incubation with PDMPO (red: chlorophyll autofluorescence, cyan: PDMPO). **(G)** Bioanalyzer cDNA profile of a single *S. setigera* cell sorted directly into the lysis buffer. **(H)** SEM image of a FIB-generated cross section of the *S. setigera* cell. va-vacuole, chl-chloroplast, cy-cytoplasm, fru-frustule. **(I)** Phylogenetic placement of the *S. setigera* sequence. Numbers at nodes are percentage bootstrap values (100X). Scale bar indicates substitutions per site. **(J)** Brightfield confocal microscopy image of *S. setigera* pure cultures, with **(K)** chlorophyll autofluorescence shown in red.

We first quantified cell behaviour such as sinking speed, highlighting heterogeneity in sedimentation rates among individual live phytoplankton cells (**Figure 5 D and E**). In parallel, we tracked biosilicification in growing and newly divided cells by incubating live specimens with a fluorescently labelled substrate and tracked its incorporation into the silica cell wall using time lapse imaging (**Figure 5 F**). On the same day, we also sorted and preserved cells for single cell transcriptomics (**Figure 5 G**) and electron microscopy (**Figure 5 H**), both approaches for which immediate, rapid, and state of the art sample processing is mandatory. Finally, cells from the same population were isolated for single cell genotyping to phylogenetically place the organism and confirm it as *Sundstroemia setigera* (**Figure 5 I**), characterize its host-associated microbiome^74^, and establish long-term cultures for further mechanistic studies (**Figure 5 J and K**).

This synergistic strategy enabled the simultaneous acquisition of sophisticated multi-modal datasets targeting a single environmental species, spanning live and subcellular imaging, single organism approaches and long term cultivation. Integrating such data will enable linking cell behaviour to subcellular structures, while single-organism techniques provide a bridge between genotype and phenotype and facilitate exploration of cell-cell interactions. Taken together, this opens up new opportunities to characterize environmental species with a level of resolution previously reserved for model systems in laboratory settings, advancing our molecular and cellular understanding of life in context.

### Cultivation of environmental organisms

The sequencing-based identification and phylogenetic placement of environmental protists is rapidly progressing ^17^, however, cultivation of environmental protists is essential for addressing gaps in understanding their diversity and ecological roles ^76^. The close proximity of the AML to sampling sites provides an opportunity to rapidly incubate sensitive environmental samples. Access to a range of incubators and microscopy allows for the identification and isolation of organisms based on morphology and size, and the optimization of growth conditions. As an example, we successfully isolated and identified 96 environmental fungal species from various locations along the European coastline (**Supplementary Table 2**). This collection of living cells, available upon request, highlights the advantages of integrating proximal laboratory facilities with sampling sites to enhance the cultivation and study of diverse protist species.

### Fostering scientific collaboration and public engagement

We deployed the AML at universities and marine stations, providing a fertile ground for strong scientific collaborations between the members of the expedition and local scientists. Our technologies and expertise were made available to more than 70 local scientists, supporting 42 projects at the host institutes ^77–80^.

During the TREC expedition, over 1,000 visitors, including school children and seniors, participated in interactive experiences centered around the AML and led by our public engagement team, the AML staff, and local volunteers. These activities made complex scientific concepts accessible through games and demonstrations in native languages ^81–83^. Furthermore, the unique lab environment encouraged the participating scientists to creatively communicate their work, through audio, visual or tactile art projects ^84,85^, blending science and art to deepen public understanding and appreciation of marine ecosystems (**Supplementary Figure 3**).

### Conclusion and outlook

The diversity of systems and workflows the AML brings to the environment, allows for a more comprehensive understanding of biodiversity, behavior, and subcellular processes in their native habitats **(Figure 5)**. This approach is complemented by environmental metadata and field samples preserved specifically for omics analyses. Integrating these diverse datasets and methodologies facilitates linking an organism’s genotype to its phenotype, and thereby its life cycle, function and impact in the environment. Connecting the images and taxonomic datasets collected during the TREC expedition with the extensive sequencing data available (e.g. metagenomics, eDNA) will enable the interpretation of the cell biology in a broader environmental context. While during the TREC expedition, there was a strong focus on marine plankton research, the modular and adaptable design of the facility ensures that it can be used for expeditions and research across a broad range of habitats.

The AML reverses the typical paradigm of a core facility by bringing the technology to the user and the sample. The AML has now joined the pool of EMBL’s core facilities and in line with EMBL’s mission, it is open for access to all scientists (https://www.embl.org/groups/mobile-labs/). In the spirit of open access, we offer to share design considerations and lessons learned from the construction and deployment of the AML with anyone interested in establishing a similar facility.

The AML has strong roots in marine biology, but its true potential lies in serving as a powerful tool for science at large. The AML should be seen as a versatile facility, with an instrument portfolio that is expected to evolve with time, adapting to its user needs and scientific questions. Future implementation could include on-site sequencing, automation of cultivation workflows or analytical methods to measure key environmental factors and adapt the sampling strategies directly in the field. By supporting researchers with advanced molecular biology and imaging tools, it can serve as a bridge between lab and field research, opening new types of research avenues and promoting interdisciplinary research and collaborations.

## Acknowledgements

We thank the TREC Consortium members, the TREC Expedition team, the TREC core partners (EMBL, the Tara Europa team, the Tara Ocean Foundation, EMBRC), and the Transversal Theme Planetary Biology Co-chairs D. Arendt, P. Bork, and R. Pepperkok.

We are grateful to the Station Biologique de Roscoff, France, the Tallinn University of Technology, Estonia, the Kristineberg Center for Marine Research and Innovation, Sweden, the Plentziako Itsas Estazioa, Spain, the Centro Interdisciplinar de Investigação Marinha e Ambiental (CIIMAR), Portugal, the Institut de Ciències del Mar (ICM-CSIC) Spain, the Stazione Zoologica Anton Dohrn, Italy and the Hellenic Centre for Marine Research, Greece for their hospitality and support during the TREC expedition. We thank the captains and crew of the research vessels that supported the field work, and the colleagues from the Roscoff Culture collection and the Basque Microalgae Culture Collection. We thank F. Neveu and M. Asselin for their enthusiasm and collaboration in the building of the AML, Monique for her unwavering support and inspiration, and S. Verstraeten, M. Major, C. Tambley, V. Meier, S. Kandels, J. Bilic Zimmermann, J. Hellgoth, E. Klose, A. Milberger and L. Burger whose essential contributions made this project possible.

We are grateful for the generous support by Zeiss, Eppendorf, Ted Pella Inc, Thermo Fisher and the Manfred Lautenschläger Foundation.

We acknowledge the access and services provided by the Imaging Centre at the European Molecular Biology Laboratory (EMBL IC), generously supported by the Boehringer Ingelheim Foundation, We thank EMBLs Electron Microscopy Core Facility (EMCF), the Advanced Light Microscopy Facility (ALMF), the Genomics Core Facility, the EMBL Beamline P14 on the PETRA III synchrotron, DESY, Hamburg and the Mobile Laboratories Core Facility of the European Molecular Biology Laboratory, generously supported by the Klaus Tschira Foundation. We acknowledge cryo-ET instrumentation from the BioEM facility of the University of Basel, computational analysis through the sciCORE (http://scicore.unibas.ch/) scientific computing center at the University of Basel.

## Author Contributions

Conceptualization T.B., P.B., C.B., J.D., M.D., G.D., O.D., B.D.E., E.F., P.G., V.H., N.L., K.M., M.O., R.P., Y.S., F.V., W.W.

Data Curation S.B., T.B., C.B., A.P.B., M.B., C.M.B., L.B.P., P.C., F.C., M.D., E.F., B.G., C.J., N.L., M.O., M.P., L.P., C.R.R., Y.S., A.S., G.T., T.W., D.Y.

Formal Analysis T.B., C.B., M.B., P.C., J.D., M.D., E.F., S.F., N.L., M.O., L.P., C.R.R., A.S., F.V., T.W.

Funding Acquisition P.B., J.D., G.D., O.D., B.D.E., P.G., V.H., N.L., S.M., R.P., Y.S., F.V.

Investigation S.B., T.B., C.B., A.P.B., M.B., C.M.B., L.B.P., P.C., F.C., J.D., M.D., G.D., O.D., E.D., B.D.E., E.F., S.F., B.G., P.G., V.H., M.J., C.J., M.L., N.L., F.M., K.M., K.M., M.O., M.P., L.P., E.W.P., A.R.R., I. R.-B., P.R., H.R., C.S.D., Y.S., H.S., A.M.S., A.S., G.T., F.V., T.W., W.W., D.Y., S.Z.

Methodology S.B., T.B., C.B., A.P.B., M.B., C.M.B., L.B.P., P.C., F.C., J.D., M.D., G.D., O.D., E.D., B.D.E., E.F., S.F., B.G., P.G., A.H., V.H., M.J., C.J., M.L., N.L., F.M., K.M., K.M., M.O., M.P., L.P., M.P., E.W.P., A.R.R., C.R.R., I. R.-B., P.R., H.R., C.S.D., Y.S., H.S., A.M.S., A.S., G.T., F.V., T.W., W.W., D.Y., S.Z.

Project Administration P.B., M.D., G.D., O.D., B.D.E., P.G., V.H., N.L., S.M., R.P., Y.S., H.S., F.V.

Resources J.D., B.D.E., E.F., P.G., V.H., N.L., K.M., Y.S., F.V.

Software T.B., E.F., A.H., L.P., C.R.R.

Supervision J.D., G.D., O.D., B.D.E., P.G., V.H., N.L., S.M., M.P., Y.S., F.V.

Validation S.B., T.B., C.B., J.D., M.D., E.F., M.L., F.M., K.M., M.O., L.P., E.W.P., H.R., Y.S., H.S., A.M.S., F.V.

Visualization S.B., T.B., M.B., P.C., J.D., M.D., O.D., E.F., N.L., F.M., M.O., L.P., A.R.R., C.R.R., H.R., H.S., A.M.S., A.S., F.V., T.W.

Writing – Original Draft Preparation C.B., M.B., P.C., J.D., M.D., G.D., O.D., B.D.E., N.L., F.M., H.S., F.V., T.W.

Writing – Review & Editing C.B., M.B., P.C., J.D., M.D., G.D., O.D., B.D.E., P.G., V.H., N.L., F.M., M.P., A.R.R., H.S., F.V., T.W.

All authors have read and approved the final manuscript.

## Methods

### The AML stops of the TREC expedtition

Between March 2023 and August 2024, we were hosted for 2-6 weeks at the following institutes: the Station Biologique de Roscoff, France, the Tallinn University of Technology, Estonia, the Kristineberg Center for Marine Research and Innovation, Sweden, the Plentziako Itsas Estazioa, Spain, the Centro Interdisciplinar de Investigação Marinha e Ambiental (CIIMAR), Portugal, the Institut de Ciències del Mar (ICM-CSIC) Spain, the Stazione Zoologica Anton Dohrn, Italy and the Hellenic Centre for Marine Research, Anavyssos, Greece. Operations paused for a winter break between November 2023 and February 2024. Due to logistical constraints, the AML instrumentation was installed at local institutes for the first two stops, while the fully operational AML was deployed for the remaining six.

### Environmental sampling and sample concentration

Plankton samples were collected using nets of various sizes (5-150 µm) for 2-10 minutes, depending on the site. Upon recovery on the boat, the samples were further fractionated using metal sieves (500 µm - 40 µm) and transferred into bottles, which were kept at ambient seawater temperature in a cooler. Upon arrival at the lab, samples were transferred to an incubator at ambient seawater temperature running a 12 hours day/night cycle and processed within 4 hours of retrieval.

When necessary, cells were initially concentrated onto a 1.2 µm mixed cellulose ester (Millipore) filter using a manual vacuum filtration unit. To prevent clogging during the process, the cell suspension was periodically resuspended using a plastic Pasteur pipette. Once the volume was reduced to 2–4 mL, the cells were resuspended from the filter surface and transferred into Eppendorf tubes.

## Cell Sorting

### Sundstroemia setigera

Natural samples from 5 μm plankton net tows were pre-filtered on 200 μm sieve and diluted using 0.22 μm filtered sea water (FSW) in 50 mL tubes (Eppendorf) before analysis in COPAS Vision 500 (Union Biometrica). Sheath fluid was 0.22 μm FSW and the fluidics system operated under 0.2–2 psi. Sample overviews were acquired using measurable cell properties - extinction (optical density), time of flight and chlorophyll autofluorescence in red. Healthy *Sundstroemia* populations identified by relative length and positive chlorophyll signal before isolation using pure-no-doubles sorting mode for downstream applications. Cells were sorted into either 96-well plates twin.tec PCR Plates LoBind (Eppendorf) for single-cell sequencing, or 96-well plates Nunc MicroWell, Nunclon Delta-Treated, Flat-Bottom Microplates (Thermo Scientific) pre-filled with 200 μL F/50 growth media for cultivation. For the PDMPO assays, cells were sorted into 50 mL tubes (Eppendorf) pre-filled with 1 mL 0.22 μm FSW. For single-cell sequencing, individual *Sundstroemia* cells were sorted into 96-well plates filled with 10 µl of 200 mM lysis buffer containing 200 mM Guanidin - hydrochloride, 167 mM DTT, 1.7 mM Tris-HCL pH 7.5, 1% TritonX.

### *Dinophysis* sp

Fresh environmental samples (size range 20-200 µm) were analysed at the COPAS Vision 500 (Union Biometrica) and a gating strategy based on fluorescence parameters (autofluorescence of chlorophyll and phycoerythrin), extinction (optical density) and time of flight was used to identify *Dinophysis* sp. cells. The brightfield images generated by the CMOS camera were used to validate the sorting gate. Roughly 300 cells were sorted in 5 minutes and deposited into a pipette tip modified to hold the 3 mm, 200 µm deep type A HPF carrier (Wohlwend). Cells were isolated in purity sorting mode to avoid possible contaminations of other cells.

## Electron Microscopy

### High-Pressure Freezing (HPF) and Freeze Substitution

1.2 µl of concentrated sample was loaded into a hexadecene coated 3 mm gold-coated copper carrier (200 µm type A), sandwiched with the flat side of the type B aluminium carrier (Wohlwend) and high-pressure frozen with the EM ICE (Leica Microsystems).

The samples were freeze-substituted at EMBL’s Electron Microscopy Core Facility using the EM-AFS2 (Leica Microsystems) with 2% osmium tetroxide in glass-distilled 100% acetone. The temperature was maintained at −90°C for 38h, gradually raised to −60°C over 15h, and held steady for another 10h. Subsequently, the temperature was increased to −30°C and then brought to 0°C at the steepest possible slope. The metallic chamber holding the samples was then removed from the AFS, and acetone rinses (5 times, 15 minutes each) were performed on ice under a fume hood. The final rinse was done at room temperature.

The samples were then gradually infiltrated with Epon resin without the accelerator DMP-30. This involved 2h incubations in 30%, 50%, and 70% Epon, followed by overnight incubation in 100% Epon at room temperature. This was followed by two 3h incubations in 100% Epon containing DMP-30, and a final 24h incubation. The resin was polymerized at 60°C for 48 hours.

For the workflow following the cell sorting of *Dinophysis* cells outlined in **Figure 4**, the samples were deposited in a pipette tip modified to hold the HPF carrier **(Figure 4 C)**. The holder was centrifuged briefly at 2000g for 2 minutes. After removal of the supernatant, the carrier was released from the pipette tip, transferred to the HPF and subjected to freeze substitution and resin embedding following protocol ^86^ used in previous studies ^70,87^.

The *Rhizoloenia* cells, shown in **Figure 5**, were freeze substituted in 0.1% uranyl acetate in dry acetone. Temperature was maintained for 72 hours at −90° C, then raised to −45°C at a rate of 2°C/hour and held steady at −45°C for an additional 10 hours. The samples were rinsed thrice in acetone, and gradually infiltrated with Lowicryl HM20 resin at 8 hour intervals and while increasing the temperature to −25° C. Once at 100% resin, the resin was exchanged three more times, in 10h intervals, before samples were UV polymerized for 48h.

### Chemical fixation and Scanning EM

Concentrated cells were gently centrifuged at 1000g for 5 minutes using a swing bucket rotor. The resulting pellet was then chemically fixed with 2% Formaldehyde and 0.5% Glutaraldehyde diluted in 1X PHEM buffer (60mM PIPES, 25 mM HEPES, 10mM EGTA and 2 mM MgCl2) pH 6.9, supplemented with 10% sucrose, modified from ^88^. Fixation was performed in a microwave (PELCO BioWave) where samples were irradiated at low power (7 on/off cycles of 100W, applied vacuum of 20 Hg). Samples were then centrifuged again, the fixative removed, and the pellet was resuspended in 1% FA in 0.1M PHEM buffer and stored at 4°C until further processing.

For SEM imaging, samples were washed with 1X PHEM buffer and cells allowed to sediment at 4°C. This was followed by sequential dehydration steps in 30%, 50%, 70%, 80%, 90% and 100% acetone at 4°C. 100% acetone dehydration was done overnight, followed by another ~3 hour exchange in 100% acetone. Cells were then resuspended in minimal supernatant, followed by critical point drying (CPD300, Leica Microsystems) in VitraPOR (Robu) filter plates. Finally, the cells were distributed on conductive adhesive carbon tape on a SEM stub, gold-sputter coated (Quorum Q150RS) and imaged on the Zeiss Crossbeam 540 at an acceleration voltage of 1.5 kV and current 700pA with a SESI detector.

### Transmission EM

Resin-embedded samples were trimmed and 70 nm ultrathin sections were collected with an ultramicrotome (Leica UC7) on formvar-coated slot grids, post-stained with uranyl acetate and lead citrate and imaged at the JEOL JEM-1400Flash at 120 kV.

### X-Ray Imaging

Micro Computed Tomography of the resin block containing the COPAS-sorted *Dinophysis* population was performed at the EMBL Beamline P14 on the PETRA III synchrotron at DESY, Hamburg. The imaging was performed in a parallel beam configuration at 23.46 keV and an off-centre 360° acquisition with 3600 projections, a 0.01 s exposure time, and an isotropic voxel size of 650 nm was used. The 3D volume representing the entire resin block was reconstructed using an in-house made reconstruction pipeline ^63,89,90^, this allowed us to visualise the distribution of the *Dinophysis* cells, segment individual cells to obtain volumes and precisely locate them in the block for downstream imaging. Guided by block surface landmarks and measurements from the 3D X-ray scan **(see Supplementary Video 3)**, we could target our cell of interest for FIB-SEM.

### FIB-SEM

### *Dinophysis* workflow

The whole-cell volume was acquired at an isotropic 20 nm voxel size at the Zeiss Crossbeam 540, using the Atlas 3D nanotomography pipeline. Milling was done at 30 kV, 1.5 nA and SEM imaging at 1.5 kV accelerating voltage and 700 pA current with an ESB detector (ESB Grid 1110V). Final dwell time for acquisition was 11 µs. Due to an automated milling error, only ~60% of the cell volume could be imaged by FIB-SEM. Raw dataset was aligned using the Fiji plugin - Linear Stack Alignment with SIFT ^91^. Slices with extensive curtaining were removed from the stack for better alignment. Organelles were segmented from this FIB-SEM stack using Amira (Thermo Fisher Scientific) for the purpose of visualisation and morphometric analyses, as indicated on **Figure 4**.

### Correlative light and electron microscopy workflow of *Sundstroemia* cells

*Sundstroemia*-sorted cells embedded in Lowicryl blocks were imaged with a confocal microscope (Zeiss LSM 780 NLO) to locate and mark the location of individual cells with a laser as described in ^55,92^. The distance of the cells to the block surface was measured from a confocal stack, taking as a reference the reflection of the laser light at the block face. The resin above the cell of interest was then removed in an ultramicrotome (UC7, Leica Microsystems) with a trimming knife (Diatome) until the cell was just below the surface. At this point small lines framing the cell of interest were branded on the block face with a near-to-infrared laser. Subsequently, the tip of the block was cut and mounted on a SEM stub (Agar Scientific) with conductive resin (Chemtronics) and sputtered with gold (Quorum, Q150RS). Branded cells identified with SEM were used to define the milling area, where a thin layer (10 nm) of platinum was deposited. A trench was then opened right in front of the predicted location of the cell, based on the measurements taken at the confocal microscope. 2D-high-resolution images of a longitudinal section of the cell of interest were then acquired with the ESB detector at 1.5 kV and 700pA.

### Cryo-ET

A concentrated cell fraction, ranging from 10 to 20 µm, was assessed for cell diversity and concentration using a light microscope prior to plunge freezing.

After glow discharging of the grids (PELCO easiGlow) the samples were vitrified. 4 µL of cell suspension was applied on 200-mesh R2/1 carbon-film coated copper grids (Quantifoil MicroTools) and plunged frozen using a Vitrobot Mark IV (Thermo Fisher Scientific). EM grids were clipped into Autogrid supports (Thermo Fisher Scientific) or custom-made AutoGrid cartridges modified for FIB under shallow angles and stored in sealed boxes in liquid nitrogen and loaded into Aquilos 2 FIB-SEM instruments (Thermo Fisher Scientific), where they were thinned with a Gallium ion beam as previously described ^93^. The resulting EM grids with thin lamellae were transferred to a 300 kV Titan Krios G3i microscope (Thermo Fisher Scientific) equipped with a BioQuantum post-column energy filter (Gatan) and a K3 direct electron detector (Gatan) or Titan Krios G4i with a Falcon4i direct electron detector and a Selectris energy filter for tomographic imaging. Tilt-series were obtained using SerialEM 3.8/4.1.2 software ^94^ or Tomo5 (Thermo Fisher Scientific). In all cases, tilt-series were acquired using a dose-symmetric tilt scheme ^95^, with 2° steps totalling 60/53 tilts per series. Each image was recorded in counting mode with ten frames per second. The target defocus of individual tilt-series ranged from −2 to −5 µm/−3 to −7 µm. Tilt series were acquired with a total dose of approximately 120 e^-^/Å^2^ and an image pixel size of 4.569 Å/pixel or 3.0444Å/pixel.

### Cryo-ET data processing for segmentation

Tomograms were subsequently reconstructed. Raw frames were aligned using MotionCor2 (version 1.5.0) ^96^ before dose-weighting the tilt-series ^97^ followed by manual removal of bad tilts. The resulting tilt-series (binned 4 times) were aligned in IMOD (version 4.11) ^98^ using patch tracking and were reconstructed by weighted back projection or were reconstructed with a SIRT-like filter. Additional features of interest, such as ribosomes and starch, were manually segmented using Dragonfly (version 2024.1 – Build 1597). The resulting segmentations were further refined in Dragonfly, which was also used for visualisation and movie preparation. For the first part of figure 3 Cryo-CARE (version 0.2.1) ^99^ was applied on reconstructed tomogram pairs from odd and even raw frames to enhance contrast and remove noise. All preprocessing steps were carried out within Scipion ^100^. Snapshots of denoised tomograms were captured using the IMOD 3dmod viewer. Denoised tomograms were used as input for automatic segmentation using MemBrain ^101^. The resulting segmentations were manually curated in Amira (version 2021.2). The resulting segmentation was visualized in ChimeraX ^102^.

### Cryo-ET data processing for subtomogram averaging

Cryo-electron tomography data was processed using WarpTools (v2.0.0dev29) ^103,104^. Initially, the stage tilt values in the .mdoc files were modified to correct for the pre-tilt of the lamella, which ensured that the generated tomograms were flat. The raw movies were converted from .eer to .tiff format using the RELION (v4.0.1) ^105,106^ function relion_convert_to_tiff. The raw movies were then motion-corrected and averaged, before contrast transfer function (CTF) estimation with a defocus range of −4.5 to −10 µm. Tilt series stacks were generated and manually visually inspected in order to remove low-quality images. Tilt series stacks were aligned using AreTomo2 ^107^. The defocus handedness was verified using WarpTools, before 3D-CTF corrected tomograms were reconstructed at a pixel size of 12.18 Å. For visualisation purposes only, tomograms were denoised with cryoCARE ^99^ followed by IsoNet ^108^.

Ribosomes were picked in non-denoised, 3D-CTF corrected tomograms using PyTOM template matching ^109^. A plant ribosome (EMDB-15806) ^110^ was used as the initial reference. Picked particles were visually inspected and false positives were deleted using ArtiaX ^111^. 643 particles from 2 tomograms were extracted as 3D-subtomograms. Euler angles were deleted from the .star file to reduce reference-bias from particle picking. 2D-classification in RELION (v4.0.1) (K=5, T=2, 200 iterations with VDAM algorithm) was used to remove junk particles. 3D refinement was carried out using a feature-less blob as the initial reference to reduce reference-bias. The resulting structure was clearly identifiable as a ribosome. The ribosome structure was then processed using M, where image warping (on a 1×1 grid) and particle poses were refined. The resulting structure from 389 particles reached a global resolution of 28.65 Å, according to Fourier shell correlation (FSC = 0.143). The structure was deposited in the Electron Microscopy Data Bank (EMDB) (EMD-56682). The ribosome was visualised as an isosurface representation. A lower isosurface threshold was used to visualise the neighbouring ribosomes in a polysome arrangement.

## Light Microscopy

### Cryo-fluorescence imaging

After high-pressure freezing, carriers were transferred to the cryo stage (Linkam Stage CMS196V, Linkam) and mounted on a Cryo Widefield Microscope (ZEISS Axio Imager 2, Zeiss) for inspection. Images were taken with 5x objective (EC Plan-NEOFLUAR 5x/0.16, Zeiss), a Chlorophyll A filter cube (Excitation filter BP430/24, Dichroic FT488, Emission filter BP685/50, Zeiss) and a colour camera (Axiocam 208, Zeiss).

A subset of the frozen down grids were imaged with a Zeiss LSM900 Airyscan 2 equipped with a Linkam cryostage and a 5x (C Epiplan-Apochromat NA 0.2), 10x (C Epiplan-Apochromat NA 0.4 DIC) and 100x objective (EC Epiplan-Neofluar NA 0.75 DIC). In all imaging modalities images for transmission light illumination, reflected light illumination and two fluorescent channels to detect autofluorescence were collected (488 nm and 568 nm). After acquiring overviews with the 5x objectives, the full grid was mapped with the 10x objective with a 2×2 tile scan covering a z-range of 70 µm. A maximum intensity projection was used to then select the grid squares of interest for further analysis. As a last step individual squares were mapped in detail with the 100x objective, collecting confocal z-stacks of 21.6 µm with 1.2 µm steps and a pixel size of 125 nm.

### Ultrastructure Expansion Microscopy

Concentrated cell suspensions were chemically fixed by adding formaldehyde (FA) to a final concentration of 4% by adding equal volumes of 8% FA diluted in filtered sea water. To preserve fragile structures, such as flagella, samples were not centrifuged before sample fixation. Fixative was removed by sedimenting samples at 800g for 1 minute and washing them with PBS twice. Samples were kept at 4°C in 1% FA in PBS until further processing.

Alternatively, cells were cryo-preserved by either high-pressure freezing as described above or by plunge freezing coverslips on a manual plunge freezer (Rhost LLC). Cells were left to sediment for approximately 15 minutes and loaded onto 12 mm coverslips previously coated with Poly-L-Lys and CellTak. Cells were allowed to attach to the surface for 30 minutes, and the supernatant was carefully removed and blotted away. Coverslips containing cells were rapidly plunged into liquid ethane and stored in homemade metal holders at liquid nitrogen temperature until further processing. Samples cryo-preserved either by plunge freezing or HPF were freeze-substituted into acetone containing 0.5% FA and 0.025% GA, which assists in membrane and epitope preservation. Freeze substitution and rehydration is described in more detail in ^43,47,48,112^. Briefly, HPF carriers were placed onto the acetone FA/GA mixture, frozen in liquid nitrogen prior, and a Leica AFS2 was used to maintain −90°C for at least 24h before ramping the temperature up to RT at 5°C/h. Plunge frozen coverslips were placed onto acetone FA/GA in 5 mL Eppendorf tubes, which were previously frozen at a 45° angle. Tubes were placed onto dry ice, completely covered with more dry ice, and left in a closed styrofoam box for at least 12h. Most of the dry ice was subsequently removed, leaving about 10% of the bottom of the box covered and samples allowed to slowly increase in temperature over the next 4-5 h before removing all remaining dry ice and letting the samples reach RT. Both HPF and plunge frozen samples were rehydrated by washes in 2x 100%, 2x 95%, 75%, 50% EtOH in ddH2O for 3 minutes each before moving them into PBS and keeping them at 4°C until use.

For U-ExM, all types of samples (chemical, HPF, plunge frozen) were anchored in 1% acrylamide and 0.7% FA in PBS by incubating them at 37°C ON. Cells in suspension were mounted onto Poly-L-Lys and CellTak coated coverslips. Coverslips were inverted onto droplets of monomer solution (MS, 10% (v/v) acrylamide, 19% (w/v) sodium acrylate, 0.1% N,N’-methylenbisacrylamide (v/v), in PBS) on ice, to which 0.5% (w/v) APS and 0.5% (w/v) TEMED were added just prior, initiating polymerisation. Samples were kept on ice for 5 minutes, and polymerised at 37°C for 45 minutes under humid conditions. Homogenisation was ensured by heating samples to 95°C in a denaturation buffer (50 mM Tris, 200 mM NaCl, 200 mM SDS, pH 9.0) for 1.5 h before expanding the hydrogels in three subsequent ddH2O washes. All stainings were performed on gels shrunken in PBS; the 12g10 anti-alpha-tubulin antibody (DSHB Cat# 12G10; RRID:AB_1157911) was used in 4% BSA in PBS-T (0.2% Tween-20) at 1:500 final dilution. After ON incubation at 37°C, gels were washed three times with PBS-T for 5 minutes each before incubating them with secondary antibody (Goat anti-Mouse, Alexa Fluor 488, Thermo Fisher Scientific Cat# A-11001; RRID:AB_2534069) in 4% BSA in PBS-T for 4h at 37°C together with Hoechst 33342. Membranes were labeled with BODIPY™ TR Ceramide (Thermo Fisher D7540) at 1 µM for 2h at 37°C in PBS. Cells were imaged by reexpanding gels in ddH2O, mounting them onto Poly-L-Lys-coated Ibidi glass bottom chambers, and acquiring images at a Nikon-CSU-W1 SORA 749 spinning disk confocal microscope with water immersion objectives (Apo LWD 40x 750 WI Lambda-S/1.15, 0.60 and SR P-Apochromat IR AC 60x WI/1.27). Image processing and visualization were done using ImageJ FIJI ^113^.

### Gravity Machine

A vertical tracking microscope, called the gravity machine ^114^ was used to obtain sedimentation rates. The optical setup utilized dark-field illumination, using an LED ring to illuminate the sample, a 10x LWD objective (Boli Optics), a 75 mm tube lens, and a Daheng MER2-630-60U3C Camera. For all images collected, the pixel size is 0.64 µm/pixel. Samples were loaded into wheels using the same method established in Krishnamurthy et al. Wheels were treated with 5% BSA solution overnight and rinsed three times with water. Wheels were then loaded with 0.2 µm filtered seawater from the same location that the samples were collected. 0.75 µm Latex beads were added to the wheel for the post-processing which uses particle imaging velocimetry. Lastly, the sorted population of *Sundstroemia* was added to the wheel. Once the wheel was mounted onto the microscope, and a cell of interest was located in the field of view, the DaSiam object tracking software ^115,116^, which was implemented into the Gravity Machine tracking software ^114^, was used to track the cell. Images and X,Y,Z stage coordinates were recorded at a frame rate of 10 frames per second. 11 tracks ranging from 20 seconds to 60 seconds were collected. All tracks were processed using the Gravity Machine Analysis Scripts developed in Krishnamurthy et al. These scripts use particle imaging velocimetry to extract the true 3D movements from the X,Y,Z coordinates of the stage and flow of the Latex beads in the wheel. Once processed, the tracks were plotted using the script “Plotting_Rhizo.py”. All data from these experiments can be found in the zenodo repository: https://doi.org/10.5281/zenodo.18508071. In the archive is a python script titled “Plotting_Rhizo.py”. This script can be run to recreate both “AllTracks.png” and “SedimentationRates. png”. Each track has its own folder, titled YYYYMMDD_HHMMSS after the date and time that each track was collected. Inside each track folder there is a folder containing the raw images, labeled DF1. The repository contains the raw track, unprocessed, titled “YYYYMMDD_ HHMMSS_unprocessedtrack.csv”, the processed, or corrected track post PIV titled “YYYYMMDD_HHMMSS_correctedtrack.csv”, and movie with a time-stamp and scale bar of the processed images titled ““YYYYMMDD_HHMMSS.avi”, and a plot of the track (Z (mm) vs Time (s)) titled “YYYYMMDD_HHMMSS.png”.

### Confocal LSM

Confocal images of a subfraction of each sample were acquired within 2 h post collection. Multiwell glass bottom dishes (CELLVIEW CELL CULTURE SLIDE, PS, 75/25 MM, 543078, Greiner bio-one) were coated with Poly-D-Lysine (A-003-M, Sigma) for 5 minutes, which captures settling organisms in place, while keeping their structure intact. Cells were concentrated as described above to achieve a dense distribution but avoiding overlapping cells, and 200 µl of each sample were transferred into a well. Live samples were stained with Hoechst 33324 (Thermo Fischer, 10 µM final concentration) to label microbial DNA and surface structures of dinoflagellates simultaneously. Automated imaging of the whole well was performed on an inverted laser-scanning confocal microscope (LSM900, Zeiss) with a 20x objective (Plan-APOCHROMAT 20x/0,8, Zeiss) with detection of Chlorophyll a (excitation with 640 nm and detection between 656–700 nm) and phycoerythrin (excitation with 560 nm and detection between 500–645 nm) autofluorescence, Hoechst33342 staining (excitation with 405 nm and detection between 410–470 nm) and transmitted light (T-PMT). Microscope data were processed into single-object patches using Serving Vision to Living Things ^117^, a Python library that provides image analysis as a network service. In short, for each multiposition .CZI microscope file, the sample metadata was retrieved, image frames were arranged in a standard channel order, then segmented in a pipeline with multiple ilastik models ^118^. Single objects were cropped and monochrome images for each channel combined with a composite were exported and uploaded to EcoTaxa via its API. For detailed steps see https://doi.org/10.5281/zenodo.18593385.

### Feedback microscopy

The adaptive feedback microscopy ^35,36^ was implemented with the AutoMicTools package (https://git.embl.org/halavaty/AutoMicTools) in Fiji ^113^ and a macro in the Open Application Development (OAD) environment of the Zeiss ZenBlue software (https://git.embl.org/grp-almf/zeiss_zenblue_automation). The OAD macro controlled acquisition of low resolution overview images and high-resolution 3D image stacks. A custom image analysis script for detecting organisms containing phycoerythrin pigment was set up in Fiji and executed within AutoMicTools for the online analysis of low resolution images. The positions of objects identified by AutoMicTools were automatically sent to the OAD macro to trigger high-resolution acquisitions of target objects.

### PDMPO assays

A pure *Sundstroemia* population was sorted into a 50 mL falcon tube using a COPAS Vision 500 (Union Biometrica). The population was incubated with PDMPO (LysoSensor Yellow/Blue DND-160, Invitrogen) at a final concentration of 0.125 μM at 12°C with a 12h day/night cycle and no added growth media. After 24h and 72h, aliquots from 50 mL falcon tubes were imaged in a Nunc Lab-Tek 8-well chambered Coverglass (Thermo Scientific) using a LSM 900 laser-scanning confocal microscope (Zeiss) and 40x objective (Zeiss Plan-APOCHROMAT 40x/1,3) with 405 nm excitation of PDMPO and 488 nm excitation of chlorophyll autofluorescence. Maximum Intensity Projections were generated from z-stacks with Fiji ^113^.

### *Sundstroemia* cultures

Cultures were started from single cells of *Sundstroemia* isolated from natural samples in Roscoff using a COPAS Vision 500 (Union Biometrica) and identified as *Sundstroemia setigera* through PCR (see MM “Cell Sorting and “Single Cell Metabarcoding”). Cultures are maintained in silica-enriched K/2 growth media at 15℃ under day/ night cycle of light at 50 μE. Cultures were imaged in an Imaging Plate 96 CG (Zell-kontakt GmbH) using a LSM 900 laser-scanning confocal microscope (Zeiss) and Zeiss Plan-Apochromat 10x/0.45NA M27-Air objective with 488 nm excitation of chlorophyll autofluorescence.

## Sequencing

### *Sundstroemia setigera* cultures DNA extraction

1.5 ml of *Sundstroemia* cultures were harvested by centrifugation and the pellet was used as starting material. DNA was extracted using DNA NucleoSpin Plant II kit (MACHEREY-NAGEL). Species were identified by PCR targeting 18S rDNA using universal eukaryotic forward primer Euk-A (AACCTGGTTGATCCTGCCAGT) and universal eukaryotic reverse primer Euk-B (TGATCCTCCTGCAGGTTCACCTAC) ^119^. The PCR was done using a Pfu-Sso7d polymerase master mix ^120^ which contained all the needed components (20 mM Tris pH 8.8, 2mM MgCl2, 60 mM KCl, 10 mM (NH4)2SO4, 0.01 mM EDTA, 0.1 % TritonX-100, 4 % Glycerol, 0,0025% Xylene Cyanol FF, 0,025 % Orange G, 0.2 mM dNTPs, 0,02 U/μL Polymerase). Both forward and reverse primers were used at a concentration of 0. 5 µM final concentration. The PCR was conducted as follows: initial denaturation step at 98°C for 30 seconds followed by 30x amplification cycles of denaturation at 98°C for 10 seconds, annealing at 67°C for 30 seconds and elongation at 72°C for 1 minute. After the amplification cycles a final elongation at 72°C for 10 minutes was performed. Amplified DNA was stored at −20°C. PCR products were purified using QIAquick PCR Purification, (Qiagen) and sent for sanger sequencing to Eurofins Genomics using PCR primers. Results were uploaded on Benchling (www.benchling.com), where forward and reverse reads were concatenated. The final sequence was deposited on Genbank with the accession number PX488268.

### Phylogeny

Representative reference sequences from each major diatom group were downloaded from NCBI, including outgroups. A total of 26 sequences were included in the analysis. Sequences were aligned using MAFFT v7 ^121^ online default parameters. The alignment was manually inspected and trimmed in Gblocks ^122^ to remove poorly aligned regions and gaps, resulting in a final alignment of 1641 positions. Maximum likelihood phylogenetic inference was performed using IQ-TREE online under the GTR+FO substitution model ^123^. Branch support was assessed using standard bootstrap with 100 replicates. The resulting tree was visualized using phylo.io ^124^.

### *Sundstroemia s*ingle cell cDNA profile

Plates were stored at −20°C on the AML, then −80°C once shipped to EMBL. cDNA synthesis and amplification was based on ^125,126^, with minor improvements from EMBLs GeneCore Facility as follows. A first step of denaturation was performed with 2.4 µl/well mixed with 1 µl of 10 mM dNTP mix and 1 µl of 5 µM oligo dT (Sigma Aldrich: Oligo-dT30VN) on ice. This mix was incubated for 3 minutes at 72°C on a pre-heated block and placed on ice directly after for at least 2 minutes. A second step of RT reaction was performed using the following mix per sample: 2 µl of SSRT IV 5 buffer (Invitrogen/ Thermo Fisher Scientific); 0.5 µl of 100 mM DTT; 2 µl of 5 M betaine; 0.1 µl of 1 M MgCl2; 0.25 µl of 40 U/µl RNAse inhibitor; 0.25 µl of SSRT IV; 0.1 µl of 100 µM TSO and 1.15 µl of nuclease free water. The RT master mix was added to the cells followed by pulse vortexing and then placed in a PCR cycler at 52°C for 15 minutes, followed by 80°C for 10 minutes and stored at 10°C. The third step of amplification was done immediately after RT reaction: 10 µl of the cDNA mix was mixed with 12.5 µl of 2x KAPA HIFI HotStart Ready Mix and 2.5 µl of nuclease free water per sample. The PCR was conducted as follows: initial denaturation step at 98°C for 3 minutes followed by 18x amplification cycles of denaturation at 98°C for 20 seconds, annealing at 67°C for 15 seconds and elongation at 72°C for 6 minutes. After the amplification cycles a final elongation at 72°C for 5 minutes was performed and samples were held at 10°C. cDNA was purified using 0.6x volume of SPRI beads (Beckman Coulter). The mixture was incubated for 5 minutes at room temperature, then placed on a magnet to separate the beads. Supernatant was completely removed and beads were eluted in 1 µl of nuclease-free water. 12 µl of the supernatant were taken out and checked on a HS DNA Bioanalyzer chip.

## Cultivation

### Environmental fungal isolation and 18S rRNA identification

Environmental samples collected from various sites were diluted 1:200 in filtered seawater and plated (200 µl) onto agar plates prepared using Marine Broth (MB; Millipore, Sigma-Aldrich, 76448) or Chromosphaera perkinsii (Cperk) medium^44,45^, supplemented with 2% agar, 200 µg/l thiamine (Sigma-Aldrich, T1270), and one of two antibiotic combinations: 100 mg/l streptomycin with 105 mg/l doxycycline (Sigma-Aldrich, S9137; Thermo Scientific, 446060250) or 50 mg/l kanamycin with 500 µg/l cloxacillin (AppliChem, A1493; Cayman Chemical, 22249). Plates were incubated at 4–17°C in complete darkness until colonies developed. Non-bacterial colonies were identified microscopically, transferred to fresh plates with or without antibiotics, and cultured until sufficient biomass was obtained for DNA extraction or colony PCR. For genomic DNA extraction, colony material was suspended in PBS containing Lysing Matrix C beads (MP Biomedicals, 6910-100), lysed using a MagNA Lyser Instrument (Roche, 05401827001; 2 cycles of 30 seconds at 4500rpm at 4°C), and processed with the QIAamp DNA Blood Mini Kit (QIAGEN, 51104) according to the manufacturer’s instructions. Extracted DNA was stored at −20 °C for further use.

For colony PCR, a small fraction of the colony was resuspended in 30 µl of 1× TE buffer, boiled at 95 °C for 10 minutes, and 1 µl of the boiled sample was used as the DNA template. The PCR reaction (20 µl total volume) consisted of 10 µl OneTaq 2× Master Mix, 2 µl each of forward primer (42F: 5′-CTC AAR GAY TAA GCC ATG CA-3′ or 82F: 5′-GAA ACT GCG AAT GGC TC-3′) and reverse primer (1498R: 5′-CAC CTA CGG AAA CCT TGT TA-3′ or 1520R: 5′-CYG CAG GTT CAC CTA C-3′), with the remaining volume made up with nuclease-free water. The PCR program included an initial denaturation at 94°C for 5 minutes (for colony PCR) or 30 s (for genomic DNA PCR), followed by 30 cycles of denaturation at 94°C for 30 seconds, annealing at 55°C for 30 seconds, and extension at 68°C for 2 minutes, with a final extension at 68°C for 5 minutes. PCR products, typically 1.000–1.800 bp, were visualized on a 1% agarose gel, purified using the Wizard SV Gel and PCR Clean-Up System, and subsequently sequenced.

